# An engineered biosensor for the fast and accurate detection of terephthalate

**DOI:** 10.64898/2026.04.03.716257

**Authors:** Marc Scherer, Philipp Wenger, Andreas Gagsteiger, Onur Turak, Birte Höcker

## Abstract

Accelerating the development of enzymatic degradation of polyesters such as poly(ethylene terephthalate) (PET) and poly(butylene terephthalate) (PBT) requires a rapid and parallelizable detection method. We developed a protein-based biosensor for the fast and accurate quantification of the PET and PBT degradation product, terephthalate (TPA), which we named TPAsense. Engineering TPAsense required overcoming low thermal stability and aggregation of the initial construct by introducing stabilizing mutations without disrupting the binding affinity to TPA. The sensor performance was validated by screening for the PBT degrading activity of a Leaf-branch Compost Cutinase (LCC) mutant library and comparing with liquid chromatography data. TPAsense detects nanomolar concentrations of TPA enabling shorter incubation times for screening workflows. In addition, a comparative analysis of PETase and PBTase kinetics was performed with TPAsense. Finally, we demonstrated the detection of PET microplastic in samples from a wastewater treatment plant by combining the biosensor and a PETase. TPAsense offers a platform to accelerate PETase and PBTase development for plastic waste recycling and detection of microplastic in the environment.

## Main text

Only a small portion of the constantly growing amount of post-consumer plastics are recycled while the rest is incinerated or disposed of in the environment ^1–4^. The conversion of plastic waste into high-value chemicals ^5,6^ promises to overcome certain limitations of current recycling strategies ^7^ and to increase circularity and economic viability. Highly efficient (bio)catalysts are required to break down plastics at mild conditions in order to reduce the demand for energy and toxic chemicals ^8,9^. Until now, only a few plastics such as poly(ethylene terephthalate) (PET), poly(butylene terephthalate) (PBT) and more recently polyurethanes are amenable to enzymatic degradation ^10–12^, but the turn-over rates are low especially in the case of PBT ^13,14^. Therefore, it is crucial to optimize existing and identify novel enzymes for plastic degradation. Although a variety of methods were developed for screening or characterizing enzymes with PET- or PBT-degrading activities ^15–22^, major limitations are the trade-off between sensitivity and throughput of screening assays and the costs of specialized equipment. Additionally, claims for standardization of PETase research methodology were raised to ensure comparability of data ^23^. Thus, having a reliable and fast detection method for TPA to identify and characterize the best PETase or PBTase variants without the need for specialized equipment would yield considerable benefits. We set out to develop a biosensor-based assay with low operational costs and short preparation times that 1) can accurately quantify nanomolar concentrations of TPA even in complex samples such as wastewater, 2) is amenable to medium-throughput screening of PETases or PBTases, and 3) allows follow-up characterization e.g. the determination of kinetic parameters.

Engineering of a biosensor requires connecting small-molecule sensing to a signal output detectable by standard laboratory equipment. Fusing periplasmic binding proteins (PBP) and a circular permuted GFP (cpGFP) to yield fluorescence-based, genetically encoded biosensors for small molecule detection has shown versatile application primarily in biomedical research ^24–26^. In PBP-based biosensors, binding of a small-molecule ligand to the PBP domain (sensing) triggers a fluorescence response of the cpGFP (signal output) through direct allosteric coupling ^27^. To achieve allosteric coupling between PBP and cpGFP, a workflow was described that comprises three main steps: 1) identification of insertion sites on the PBP structure for cpGFP fusion, 2) optimization of linkers between both parts and 3) if necessary, random mutagenesis and screening to further improve sensor parameters ^27^. A PBP called TphC binds TPA with an affinity of ∼ 400 nM ^28^ and served as the starting point for this work.

### Engineering TPAsense required stability optimization

To identify potential insertion sites for cpGFP, changes in backbone dihedrals between the open and closed state of TphC (PDB IDs: 7NDS & 7NDR) were calculated (Fig. S1a) ^29^. We reasoned that positions showing large sign changes in C_ɑ_ dihedrals of neighbouring residue pairs might generate a strong allosteric coupling between TphC and cpGFP when fused together at these sites. Two residue pairs, P61-G62 & K177-G178, were identified that are located close to the ligand binding interface (Fig. S1b). A third insertion site located on an unstructured part of the hinge region close to the C-terminus was chosen (residue V283, Fig. S1b) using knowledge of a previously engineered biosensor for agmatine ^30^. We fused TphC to superfolder cpGFP (sf-cpGFP) at these three insertion sites using proline-proline linkers between both parts (Table S1).

The TphC-sf-cpGFP fusions were expressed in *E. coli* and screened for activity in lysate. Out of the three constructs, only TphC-iV283-sf-cpGFP showed a TPA-dependent change in sf-cpGFP fluorescence and was named TPAsense1 (Fig. 1a, first panel). The fluorescence intensity change (*ΔF*) of TPAsense1 upon increasing concentrations of TPA was confirmed by *in vitro* titrations with a dynamic range *ΔF/F* (*ΔF/F* = (*F_max_* − *F_min_*)/ *F_min_*) of -0.45 (Fig. 1b - first panel, Table 1). TPAsense1 was properly structured as judged by CD spectroscopy and showed a single monodisperse peak in analytical size exclusion chromatography (SEC) (Fig. S2). Nevertheless, TPAsense1 aggregated during isothermal titration calorimetry (ITC) measurements (data not shown). This observation was in line with a reduced *T_M_* of the TphC part of 43 °C upon sf-cpGFP insertion (Fig. 1c - first panel) compared to 61 °C of TphC alone ^31^. The tendency of TPAsense1 to aggregate made it necessary to optimize its thermal stability.

**Fig. 1.**
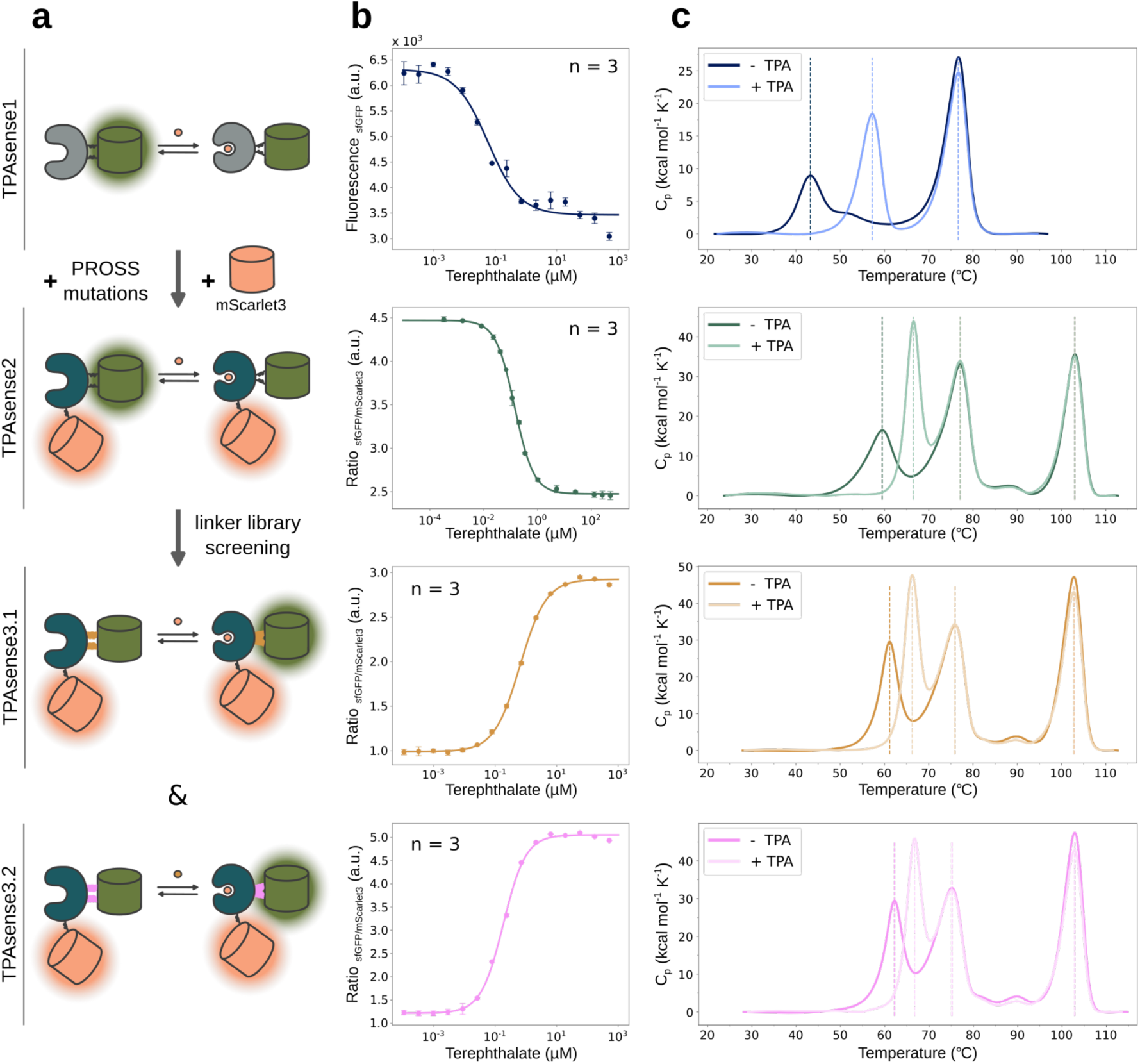
Engineering and design of TPAsense. **a**, The engineering and design workflow started with screening of potential insertion sites leading to the identification of TPAsense1. Next, PROSS mutations were introduced into TphC and mScarlet3 was added to the N-terminus for signal normalization to generate TPAsense2. A random linker library was screened to identify the final biosensor variants TPAsense3.1 and TPAsense3.2. **b**, *In vitro* titrations of TPAsense variants with serial dilutions of TPA. The data points and error bars reflect the average and standard deviations of three independent technical repeats, respectively. **c**, Thermal unfolding graphs obtained by differential scanning calorimetry (DSC) with and without TPA. Due to the shift in the unfolding temperature of *T_M_* 1, this peak is assigned to the TphC part of the TPAsense variants. As mScarlet3 is not present in TPAsense1, the second peak can be assigned to sf-cpGFP (*T_M_* 2). Consequently, the third peak only present in TPAsense2 and TPAsense3 variants is assigned to mScarlet3 (*T_M_* 3).

**Table 1.**
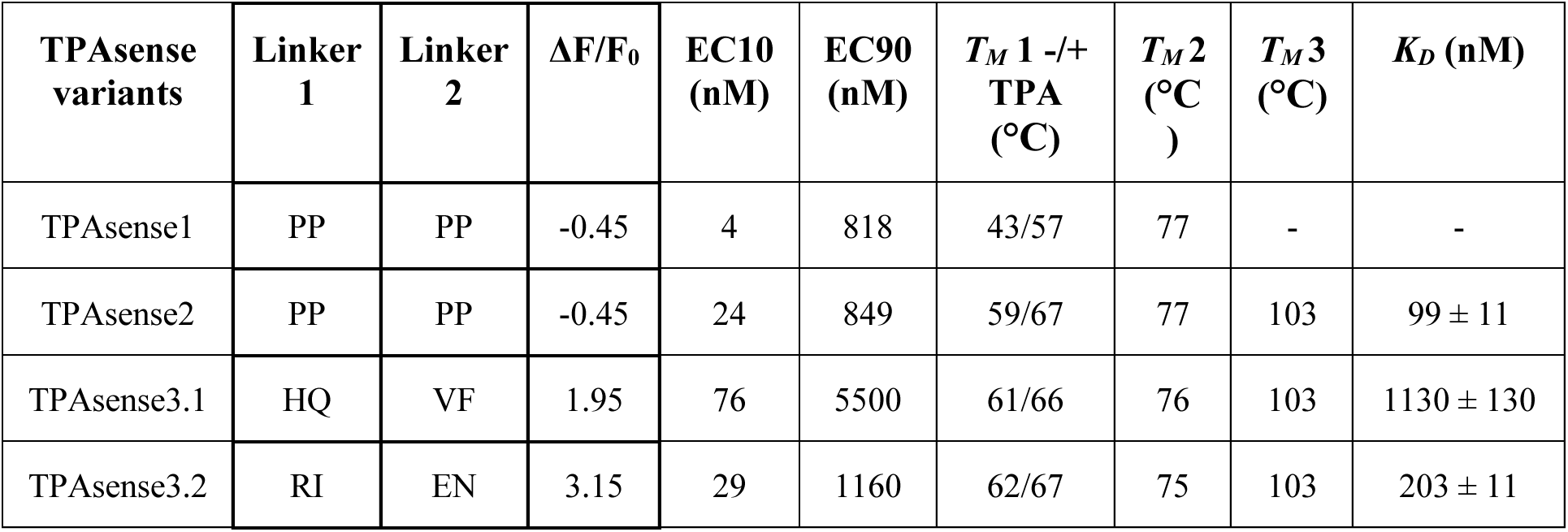
Summary of the TPAsense variants. Linker sequences are reported in single letter amino acid code. The sensor parameters are obtained from *in vitro* titrations (ΔF/F_0_ and EC10-EC90), DSC (*T_M_*) and ITC measurements (*K_D_*). The *K_D_* values reflect the average and standard deviation of three independent technical repeats.

| **TPAsense variants** | **Linker 1** | **Linker 2** | **$\Delta F/F_0$** | **EC10 (nM)** | **EC90 (nM)** | **$T_M$ 1 -/+ TPA (°C)** | **$T_M$ 2 (°C)** | **$T_M$ 3 (°C)** | **$K_D$ (nM)** |
| --- | --- | --- | --- | --- | --- | --- | --- | --- | --- |
| TPAsense1 | PP | PP | -0.45 | 4 | 818 | 43/57 | 77 | - | - |
| TPAsense2 | PP | PP | -0.45 | 24 | 849 | 59/67 | 77 | 103 | 99 ± 11 |
| TPAsense3.1 | HQ | VF | 1.95 | 76 | 5500 | 61/66 | 76 | 103 | 1130 ± 130 |
| TPAsense3.2 | RI | EN | 3.15 | 29 | 1160 | 62/67 | 75 | 103 | 203 ± 11 |

We introduced mutations into the TphC part of TPAsense1 using a PROSS (protein repair one-stop shop) design method that was developed to increase the thermal stability of PBPs without reducing the affinity to their ligands ^31^. In addition, we added a second fluorescent protein, mScarlet3, to the N-terminus of TphC for signal correction ^25^. The resulting variant, TPAsense2 (Fig. 1a - second panel), kept the same *ΔF/F_0_* of -0.45 (the fluorescence ratio of sf-cpGFP and mScarlet3 was used as output signal) but exhibited lower signal deviations between repeats (Fig. 1b - second panel) and an increased *T_M_* of the TphC part of 59 °C (Fig. 1c - second panel). Furthermore, ITC experiments confirmed comparable binding affinities of 99 ± 11 nM and 282 ± 39 nM for TPAsense2 and TphC including the same stabilizing mutations ^31^ (Table 1, Table S3, Fig. S3), respectively.

Next, we screened for variants with a direct response upon TPA addition (positive *ΔF/F_0_* values) by randomizing the linker residues of TPAsense2. 570 variants were screened in lysate (Fig. S4) and two variants, TPAsense3.1 and TPAsense3.2, were selected for further characterization. *In vitro* TPA titrations revealed a ΔF/F_0_ of 1.95 and 3.15 for TPAsense3.1 and TPAsense3.2, respectively (Fig. 1b - third and fourth panel), which was sufficient for our purposes. Notably, for a given linker library size of 160,000 variants, only a very small portion of the library was screened, thus, leaving a chance that variants with higher *ΔF/F_0_* values could be identified. While the thermal stability of both TPAsense3 variants was comparable to TPAsense2 (Fig. 1c - third and fourth panel), the binding affinity of TPAsense3.1 was reduced more than 5-fold (Table 1, Table S3, Fig. S3). Interestingly, TPAsense3.1 exhibited an increased operational range EC10-EC90 (range of quantifiable effective concentrations (EC) of ligand that lead to sensor signals between 10-90 % of the signal maximum) of 76 nM - 5,500 nM compared to TPAsense3.2. Conversely, TPAsense3.2 was more sensitive, detecting as low as 24 nM TPA while having a narrower operational range. We assessed if the TPAsense3 variants selectively interact with TPA and not with PET hydrolysis intermediates, such as mono(2-hydroxyethyl)terephthalate (MHET) or bis(2-hydroxyethyl)terephthalate (BHET). Although a sensor response was observed at higher concentrations of MHET and BHET, TPA contaminations were identified in both stock solutions using ultra high performance liquid chromatography (UHPLC) indicating that the TPAsense3 variants selectively sense TPA (Fig. S5).

Based on the differences of the TPAsense3 variants, we reasoned that TPAsense3.1 might be more suitable for screening enzymatic PET or PBT degradation of mutant libraries because a larger operational range might help to distinguish between variants with similar activities. On the other hand, TPAsense3.2 was chosen for quantifying PETase kinetics and detection of microplastic where sensing of lower TPA concentrations is required.

### Fast and accurate screening of PBTase enzymes

Recent studies showed that some PET-hydrolyzing enzymes can also degrade the structurally similar plastic PBT but with slower turn-over rates underlining the necessity for further optimization ^13,14^. We tested if TPAsense3.1 can be used to accurately screen a LCC mutant library for PBT degradation activity (see Methods for details, Fig. 2a). In total, 768 PBT degradation samples were tested. Obtaining a calibration curve (Fig. 2b) allowed the conversion of the sensor signal into TPA concentration (Fig. 2c). Signal values outside the boundaries of the operational range (EC10-EC90) of TPAsense3.1 were filtered out (Fig. 2c, gray boxes). We compared the TPA quantifications of TPAsense3.1 to those from UHPLC measurements using 527 data points after filtering. The TPA concentrations obtained with TPAsense3.1 showed a strong correlation to those from UHPLC with a R^2^ of 0.92 (Fig. 2d, Fig. S6). In addition, the slope and intercept of the linear regression was determined to be 1.08 and 0, respectively, indicating that only a minor bias was introduced by TPAsense3.1. We conclude that TPAsense3.1 can accurately quantify TPA titres of enzymatic PBT degradation. The assay can be performed in approximately 10 min per 96-well plate including a 5 min incubation of TPAsense3.1 and the sample to ensure that the equilibrium is reached. Thus, we estimate an increase in throughput of screening PBTases or PETases with TPAsense3.1 by 15-fold from ∼ 300 samples per day using UHPLC to ∼ 4,600 samples per day (assuming an 8 hour workday).

**Fig. 2.**
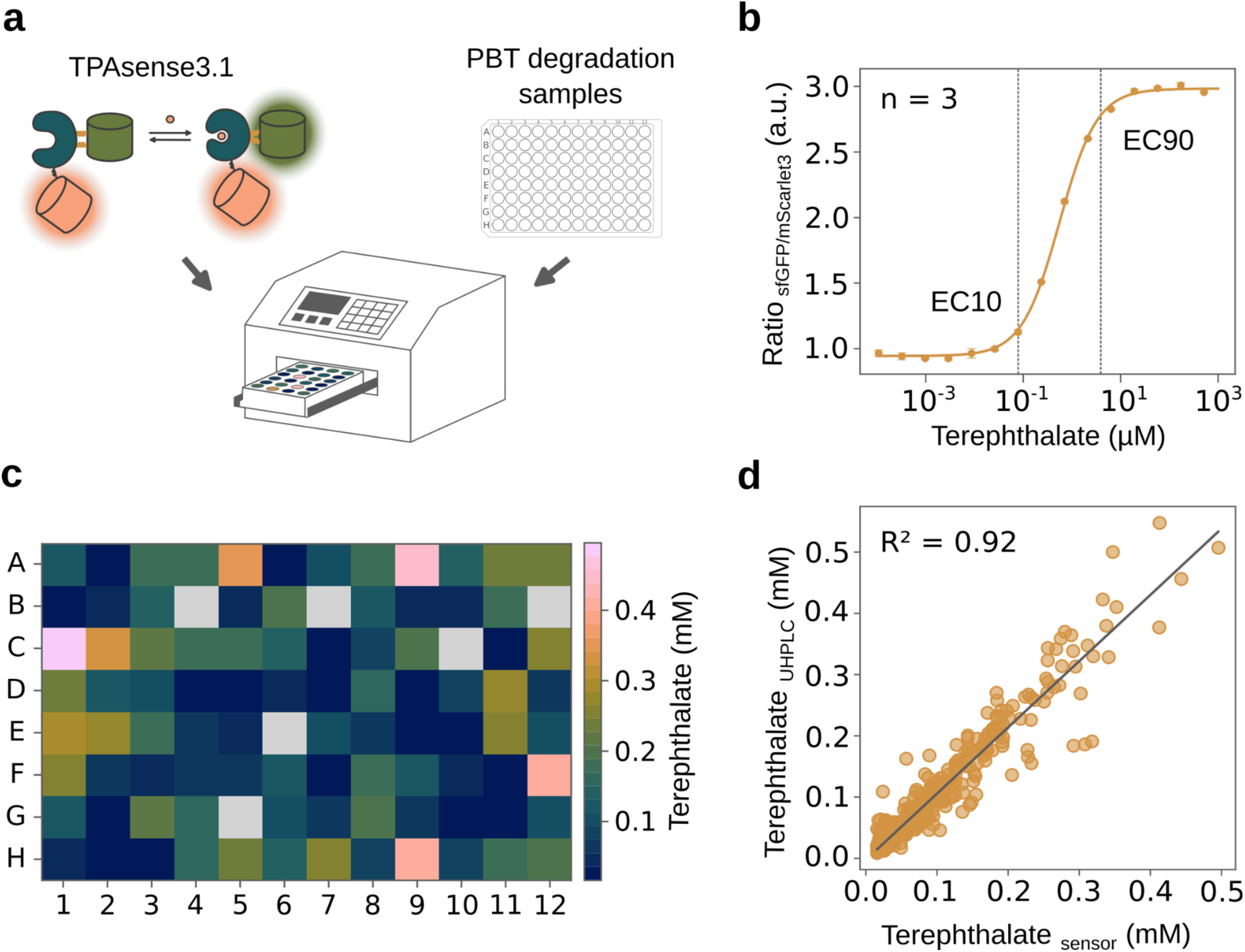
TPAsense3.1 accurately determines TPA concentrations of samples from a LCC mutant library screened for PBTase activity. **a**, The TPAsense3.1 stock solution was mixed with dilutions of the PBT degradation samples and the fluorescence signal was measured with a microplate reader. **b**, TPAsense3.1 calibration using a serial dilution of TPA and three technical repeats. The Hill equation was fitted to obtain calibration parameters. The operational range (EC10-EC90) is indicated by dashed lines. **c**, The TPA concentrations of one 96-well plate were calculated using the calibration parameters and the factor of dilution. Values outside the boundaries of the operational range of TPAsense3.1 were filtered out (gray boxes). **d**, Comparison of TPA concentrations (527 data points from 8 x 96 samples) determined with TPAsense3.1 and UHPLC. The data was fitted with a linear regression model to obtain the slope, intercept and correlation coefficient R^2^.

### Studying enzymatic PET and PBT degradation kinetics

Based on the capability of TPAsense3.1 to accurately determine TPA concentrations, we assumed that the kinetics of PETases and PBTases can be studied as well. To reduce the incubation times, we chose the more sensitive variant TPAsense3.2 for these experiments. We set out to study the effect of temperature on the reaction rates of the PETase from *Ideonella sakaiensis* (*Is*PETase) ^32^ and compare the hydrolysis rates of PET and PBT substrates with the engineered LCC variant, LCC-ICCG ^33^. First, we measured progress curves of both enzymes over 2.5 h to identify the linear phase of the hydrolysis reactions (Fig. S7). To obtain kinetic parameters of PETases and PBTases with their solid substrates, an inverse Michaelis-Menten (^inv^MM) approach ^34,35^ was employed in which the substrate load was held fixed and the product formation was followed by TPAsense3.2 as a function of the enzyme concentration. In general, the ^inv^MM model provided a satisfactory description of the experimental data (Fig. 3). In the case of PET hydrolysis using *Is*PETase, we observed a 2-fold increase of the reaction rate when the temperature was increased from 20 °C to 30 °C (Fig. 3a, Table 2). A similar acceleration of the reaction was seen from 30 °C to 40 °C but only for enzyme concentrations lower than 0.5 µM. At 50 °C, the reaction rates dropped to levels measured at 20 °C. Overall, these findings are in line with the reported temperature optimum of *Is*PETase at ∼ 40 °C ^32^. The reduced *^inv^V_max_/S_0_* (maximal reaction rate when substrate is saturated with enzyme) for *Is*PETase at 40 °C might be due to rapid reduction of available superficial ester bonds (attack sites) that can be accessed by the enzyme ^36^. In addition, bulk aggregation and denaturation of the enzyme adsorbed to the PET surface might contribute to this effect ^37^, especially at 50 °C which is above the *T_M_* of *Is*PETase ^38^. Next, we analyzed the hydrolysis kinetics of LCC-ICCG with PBT and PET as substrates (Fig. 3b, Table 2) and observed a 160-fold increase in the *^inv^V_max_/S_0_* for PET compared to PBT which is in line with the low yield reported for enzymatic PBT degradation ^13,14,39^. As the *^inv^V_max_/S_0_* is proportional to the number of attack sites ^34,35^, this analysis indicates that PET offers more accessible ester bonds to be cleaved by LCC-ICCG compared to PBT. A *^inv^V_max_/S_0_* of 21 ± 2 nmol g^-1^ s^-1^ was reported recently for LCC-ICCG at 65 °C (PET substrate) ^39^ which fits well to the *^inv^V_max_/S_0_* of 68.3 ± 12.0 nmol g^-1^ s^-1^ for LCC-ICCG at 80 °C. We conclude that the quantification of TPA with TPAsense3.2 enables the comparative kinetic analysis of PET- and PBT-hydrolyzing enzymes.

**Fig. 3.**
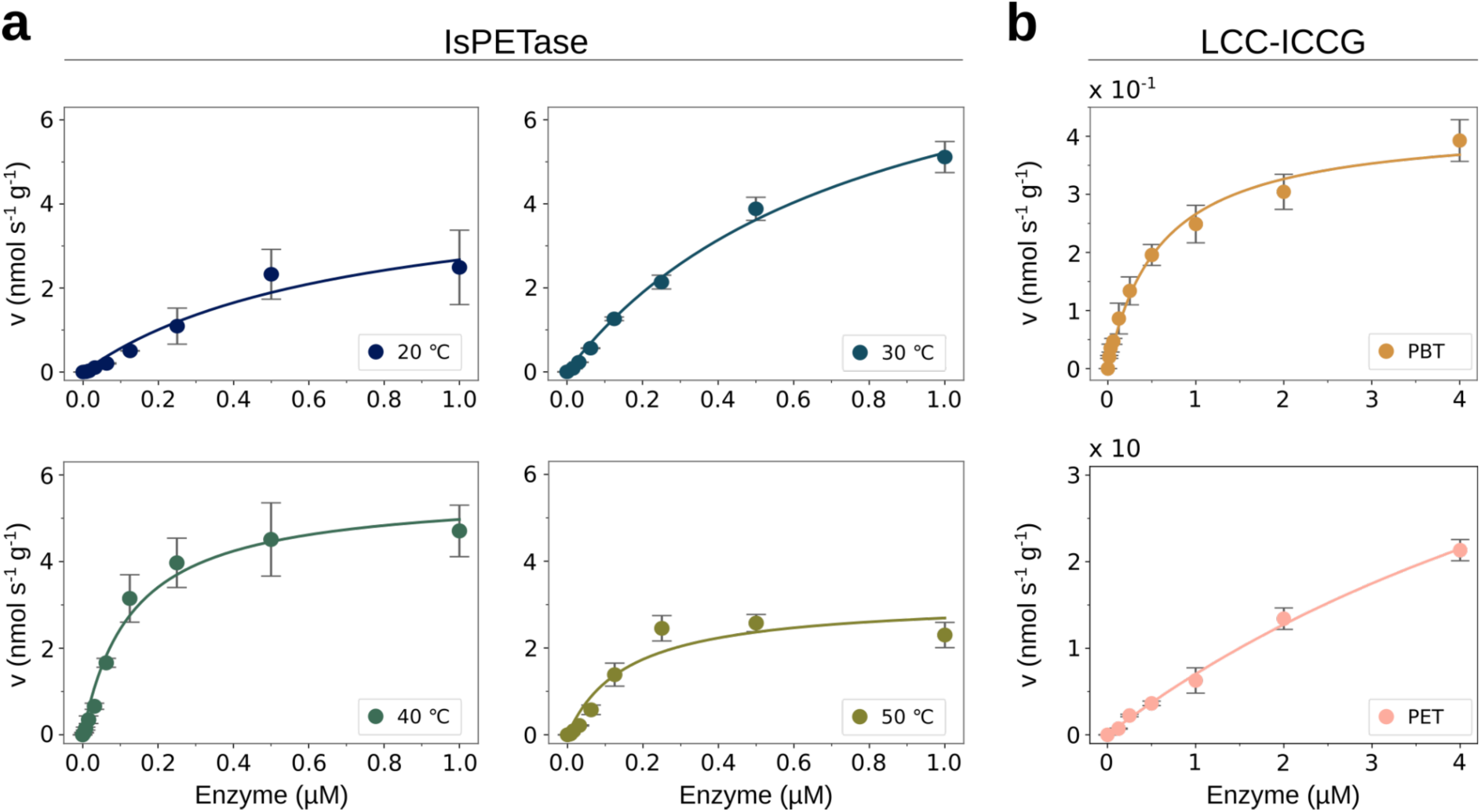
Inverse Michaelis-Menten plots of *Is*PETase and LCC-ICCG enzymes determined with TPAsense3.2. Reaction velocities were plotted in units of TPA concentration released per second and grams of polymer substrate. The data points and error bars reflect the average and standard deviations of three independent technical repeats, respectively. Lines represent the best fits of the inverse Michaelis-Menten equation (see Methods). **a**, PET hydrolysis rates as a function of *Is*PETase concentration at 20 °C, 30 °C, 40 °C and 50 °C. For each condition, the hydrolysis reaction was stopped after 2 h. **b**, PBT (upper panel) and PET (lower panel) hydrolysis rates as a function of LCC-ICCG concentration at 80 ℃. The hydrolysis reaction was stopped after 30 min for PBT and after 15 min for PET.

**Table 2.** Inverse Michaelis-Menten parameters determined for *Is*PETase and LCC-ICCG using TPAsense3.2. *^inv^K_M_* and *^inv^V_max_/S_0_* values and their standard errors were obtained from fitting the data with the inverse Michaelis-Menten equation.

| Enzyme | $^{inv}K_M$ ($\mu\text{M}$) | $^{inv}V_{max}/S_0$ ($\text{nmol s}^{-1} \text{g}^{-1}$) | Substrate | Temperature |
| --- | --- | --- | --- | --- |
| *IsPETase* | $0.70 \pm 0.30$ | $4.54 \pm 1.03$ | PET | 20 °C |
| *IsPETase* | $0.79 \pm 0.13$ | $9.32 \pm 0.84$ | PET | 30 °C |
| *IsPETase* | $0.13 \pm 0.02$ | $5.62 \pm 0.34$ | PET | 40 °C |
| *IsPETase* | $0.16 \pm 0.07$ | $3.12 \pm 0.44$ | PET | 50 °C |
| LCC-ICCG | $0.59 \pm 0.08$ | $0.42 \pm 0.02$ | PBT | 80 °C |
| LCC-ICCG | $8.72 \pm 2.07$ | $68.3 \pm 12.0$ | PET | 80 °C |

### Detection of PET microplastic in wastewater

To study the extent of PET microplastic pollution in the environment, a detection assay has to perform robustly in complex media containing a variety of different chemical compounds that might interfere with the sensor and the read-out method. Thus, microplastic detection in wastewater samples served as a challenging test case to assess proper functioning of TPAsense3.2 in complex media. We took samples from five stages of the wastewater treatment process (Fig. 4a): influent (after initial filtering of particles > 6 mm), sedimentation, denitrification, nitrification and effluent. First, we evaluated if TPAsense3.2 is capable of detecting spiked TPA in serial dilutions of each wastewater sample. TPAsense3.2 detected TPA in the same range of concentrations as the buffer control (Fig. 4b). An increased background signal was observed in influent and sedimentation samples compared to the buffer control which might be due to the higher turbidity of these samples. For the detection of PET microplastic, the wastewater samples were incubated with the PETase LCC-ICCG for 24 h at 70 °C. To test if LCC-ICCG is active in wastewater, control samples were spiked with PET microparticles (PET-MPs). All samples, except the effluent, showed substantially increased TPAsense3.2 signals compared to the negative control without LCC-ICCG or PET-MPs (Fig. 4c). The sensor response of samples spiked with PET-MPs reached the maximum signal comparable to that observed in the experiments with spiked TPA (Fig. 4b). We conclude that TPAsense3.2 can detect PET microplastic in wastewater samples.

**Fig. 4.**
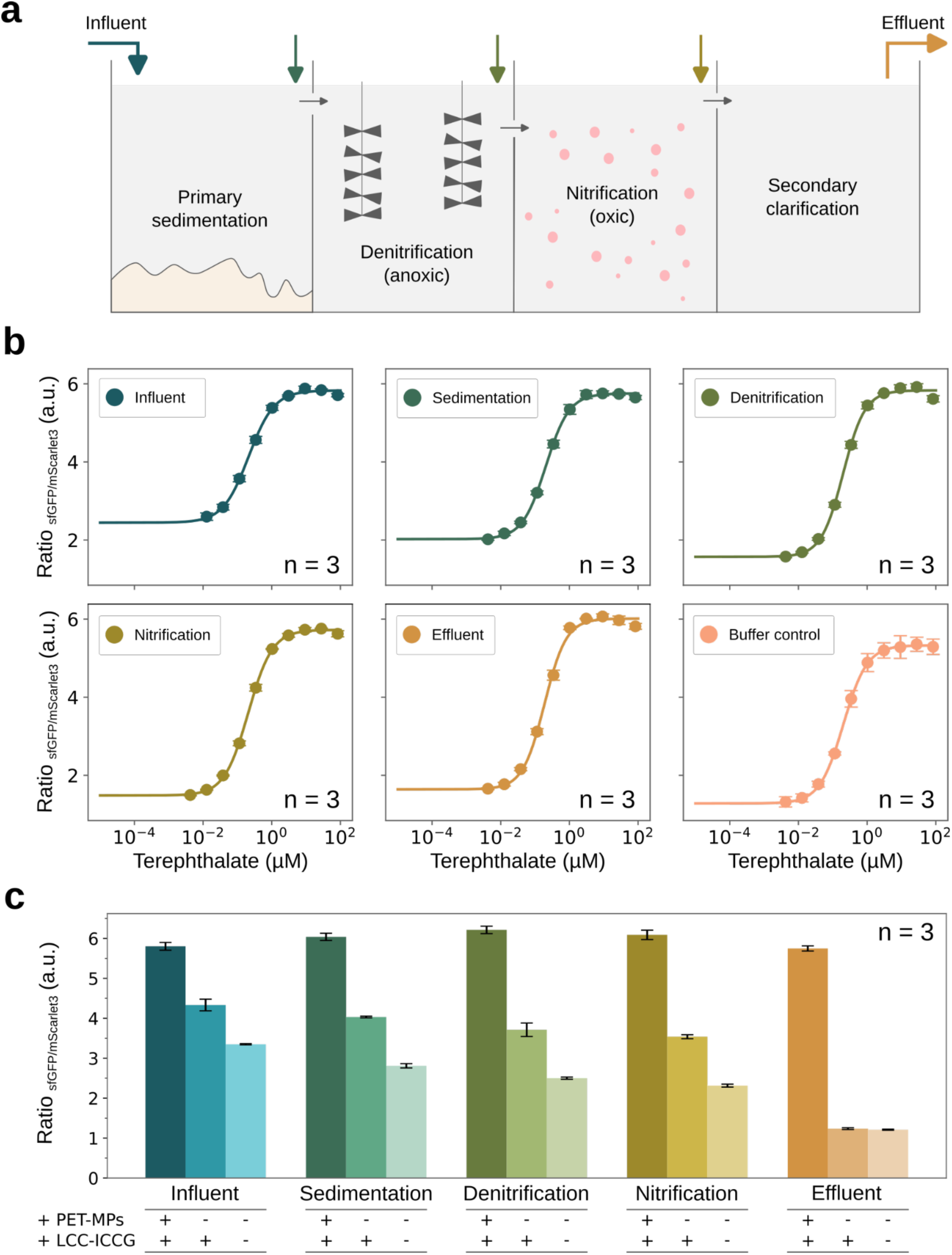
TPAsense3.2 can detect PET microplastic in wastewater samples. **a**, Schematic of a simplified wastewater treatment process. Colorful arrows indicate the approximate sampling location of influent, sedimentation, denitrification, nitrification and effluent samples. **b**, *In vitro* titrations of TPAsense3.2 with TPA-spiked wastewater samples compared to a control using the assay buffer. **c**, Responses of TPAsense3.2 to wastewater samples treated with LCC-ICCG compared to those spiked with PET-MPs and those without LCC-ICCG and PET-MPs. Each bar represents the average of three technical repeats and the error bar its standard deviation.

All samples except the effluent contained PET-MPs indicating that the removal of PET microplastic occurs in the second clarification step. As a consequence, PET microplastic should accumulate in wastewater sludge which is in line with a recent study ^40^. Still, there is evidence that some PET microplastic can be present in the effluent of wastewater treatment plants ^41^. It is possible that small amounts of PET microplastic were present in the effluent sample that accumulated to TPA concentrations lower than ∼ 30 nM (limit of quantification of TPAsense3.2).

## Conclusion

There is a necessity for standardization of PETase and PBTase research in order to properly compare results and pave the way for the transition to industrial-scale application ^23^. A reliable and fast detection method facilitating the 1) identification, 2) characterization and 3) upscaling of enzymatic PET and PBT depolymerization could support this claim for standardization. To engineer a TPA biosensor fulfilling these criteria, we had to overcome a reduced soluble expression and thermal stability of the initial TPAsense variant which is a common trade-off upon protein domain insertion ^42,43^. A reduced thermal stability of the native fold and its connection to low soluble expression ^44,45^ might affect the success rate of the insertion site screening step because potentially functional but unstable variants will not be identified. We argue that stability optimization of the sensing domain by computational design should be included into the PBP biosensor engineering pipeline as summarized by Nasu et al. ^27^ prior to insertion site screening. During the development of the biosensor assays for PETase and PBTase screening and characterization, calibration of the TPAsense3 variants using the exact condition (e.g. buffer concentrations, additives, temperature) was essential to ensure reliability of the output. We demonstrate that TPAsense3.1 accurately quantifies the TPA titres of LCC variants degrading PBT. The sensitivity of TPAsense3 variants should enable the detection of TPA in PBT degradation experiments after minutes. Utilizing this capability, we propose that PBTase screening can be accelerated by reducing the incubation time of enzyme and substrate. Additionally, the lower nanomolar limit of quantification of TPAsense3 variants might be useful in identifying enzymes with a low initial activity to PET or PBT which could then be optimized in subsequent rounds of engineering or design. The throughput of the screening assay could be further increased by using pipetting robots and an automatic injector to add the biosensor to the samples.

We showed that the fast quantification of TPA with TPAsense3.2 can be leveraged to determine kinetic parameters of PET- and PBT-hydrolyzing enzymes. Still, concerns have been raised about comparing PETase variants and reaction conditions based on initial rates as those do not necessarily reflect the product yield after extended time spans ^46^. We argue that instead of extracting ideal reaction conditions, such kinetic analysis can deepen our understanding of the mechanisms driving enzymatic depolymerization. Additionally, this approach can inform enzyme engineers on specific problems that need to be solved to achieve fast turn-over rates for example in the case of PBT hydrolysis. Taken together, we envision that TPAsense3.1 and TPAsense3.2 can simplify and speed up PETase and PBTase enzyme screening and their follow-up characterization.

A directive released by the European Union from 2024 emphasizes the necessity to monitor microplastics in municipal waste water treatment plants ^47^. Our study revealed that TPAsense3.2 is active in wastewater offering a first step towards microplastic monitoring. The subsequent experiments using the LCC-ICCG enzyme to degrade microplastics and biosensor-based detection indicated the presence of terephthalate-containing MPs in most samples of the wastewater treatment process on that particular day. Although it is likely that the detected TPA originates from PET-MPs, there is a chance that other terephthalate-containing plastics were hydrolyzed by LCC-ICCG, thus contributing to the sensor signal. To detect very low amounts of PET microplastic in samples from soil or water bodies, further engineering of TPAsense3.2 toward sub-nanomolar affinity to TPA is likely needed. Encouraged by our results, we anticipate that TPAsense3.2 can provide a means to assess the level of pollution with PET microplastic in wastewater treatment plants and other complex environments.

## Methods

### Dihedral change analysis

Open and closed state crystal structures of TphC (PDB IDs: 7NDS & 7NDR) were used as input. The dihedral of the n-th C_ɑ_ atom (C_ɑ_^n^) was calculated based on the 3D coordinates of the C_ɑ_^n-1^, C_ɑ_^n^, C_ɑ_^n+1^ and C_ɑ_^n+2^ atoms using the biopython package ^48^. The C_ɑ_ dihedral changes were then calculated by subtraction of the C_ɑ_ dihedrals of the open conformation from the C_ɑ_ dihedrals of the closed conformation.

### Computational protein design

The computational design method was described elsewhere in more detail ^31^. In short, hinge residues were identified in the closed state crystal structure of TphC (PDB ID 7NDS) using the first mode of a Gaussian network model (distance cut-off: 10 Å, spring constant: 1) ^49,50^. The PBP was split at the hinge residues to obtain two separate lobes. Using this split version of TphC, the interface and hinge residues between the lobes were identified and held fixed during the subsequent design calculations. PROSS design calculations ^51,52^ were run separately on the open and closed state crystal structures (PDB ID 7NDR & 7NDS) of TphC. Additionally, the binding pocket residues 6 Å around the ligand were held fixed. Designs with 4-10 % mutational load and the same Rosetta energy cut-offs were chosen. Finally, we removed mutations that were destabilizing in one of the two designs (two-state filter). The final design contained 9 mutations compared to the wildtype TphC protein.

### Protein expression

All genes were synthesised and cloned into a pET21b(+)-vector by BioCat GmbH. The vectors were used to transform Top10 *E.coli* cells and clones were verified by Sanger sequencing. For expression, BL21 (DE3) cells were transformed with the vectors and plated on LB agar plates containing 100 µg/mL ampicillin as selection marker. Single colonies were inoculated in 25 mL LB media-based overnight cultures supplemented with 100 µg/mL ampicillin. The main culture (0.5 L TB media) was inoculated with 5 mL of the overnight culture and incubated at 37 ℃ until the OD_600_ reached a value of ∼ 0.7. Subsequent overexpression was induced by adding isopropyl-ꞵ-thiogalactoside (IPTG) to a final concentration of 1 mM and the main culture was incubated for ∼ 18 h at 18 ℃. The cells were spun down by centrifugation (Beckman Coulter, Avanti JXN-26, JLA8.1000 rotor) at 4000 g, 4 ℃ for 20 min. After washing the cell pellets with 30 mL lysis buffer (50 mM NaH_2_PO_4_/Na_2_HPO_4_, 150 mM NaCl, 20 mM Imidazole, pH 8.0), cell suspensions were centrifuged again (Eppendorf 5920R) at 4000 g, 4 ℃ for 10 min.

### Protein purification

The cell pellets were resuspended in 30 mL lysis buffer supplemented with protease inhibitor mix HP (Serva), sonicated (Branson 6.3 mm tip, 2 x 3 min, 40 % duty cycle, output power 4) and centrifuged (Beckman Coulter, Avanti JXN-26, JA25.50 rotor) at 18 000 g, 4 ℃ for 1 h. The following purification steps were performed on an ÄKTA system. A His-Trap HP 5 mL column (GE Healthcare) equilibrated with lysis buffer was loaded with supernatant and then washed with 10 column volumes (CV) of lysis buffer before performing a stepwise elution (elution buffer: 50 mM NaH_2_PO_4_/Na_2_HPO_4_, 150 mM NaCl, 500 mM Imidazole, pH 8.0). Protein fractions were pooled and either the SUMO-protease SenP2 (initial TPAsense variants contained a SUMO-tag) or TEV-protease (TPAsense2-3 variants) was added (protease:target protein 1:10). Samples were dialyzed overnight against lysis buffer supplemented with 1 mM DTT at 4 ℃. The dialyzed samples were loaded on an equilibrated His-Trap HP 5 mL column (GE Healthcare) and washed with 10 CV of lysis buffer while the target protein was collected from the flow-through. Protein fractions were pooled and concentrated using centrifugal concentrators to a final volume of 10 mL and loaded onto a preparative size exclusion column (HiLoad Superdex 75 26/60, GE Healthcare) that was equilibrated with running buffer (50 mM NaH_2_PO_4_/Na_2_HPO_4_, 150 mM NaCl, pH 8.0). Fractions corresponding to monomeric protein were pooled and concentrated for the subsequent measurements. Protein concentrations were checked using a NanoDrop Spectrophotometer (Eppendorf BioPhotometer D30). Expressions and purifications were validated by SDS-PAGE. Samples were flashfrozen in liquid nitrogen and stored at - 80 ℃.

### Sensor activity assay in lysate

Lysate was prepared as described above (see Method: Protein purification) but instead of lysis buffer, 10 mL of activity buffer (50 mM NaH_2_PO_4_/Na_2_HPO_4_, 150 mM NaCl, pH 8.0) was used. 100 µL of lysate was mixed with 100 µL of either activity buffer or 1 mM TPA (500 µM final; dissolved in the activity buffer) in 96-well plates (black bottom Nunc A/S, Fisher Scientific). For titration experiments, serial dilutions of the TPA stock solutions were added to the lysate. After 5 min of incubation, samples were measured in a microplate reader (Spark, Tecan) with the following settings: excitation/emission with monochromator 485/535nm (sf-cpGFP) and 569/592nm (mScarlet3), bandwidth 5 nm, mirror 50 %, gain 75.

### Sensor activity assay in vitro

10 µL of 400 nM sensor protein (200 nM final) was mixed with 10 µL of either activity buffer (50 mM NaH_2_PO_4_/Na_2_HPO_4_, 150 mM NaCl, pH 8.0) or serial dilution of 1 mM TPA stock in 384-well plates (black, round-bottom, low-volume, no-bind, Corning). After 5 min of incubation, samples were measured in a microplate reader (Spark, Tecan) with the following settings: excitation/emission with monochromator 485/535nm (sf-cpGFP) and 569/592nm (mScarlet3), bandwidth 7.5 nm, mirror: Dichroic 510, gain 100 (sf-cpGFP) and 120 (mScarlet3).

For the detection of PBT degradation activity, each TPAsense3.1 stock (solutions were freshly prepared on each day of use) was calibrated by titration against serial dilutions of TPA as described above. As the UHPLC quantifications of TPA were performed beforehand, 1:200 dilutions of PBT degradation samples using the activity buffer were made to approximately reach the operational range of TPAsense3.1. If no prior knowledge of the TPA titre is available, iterative testing with different dilutions might be required for TPA quantification. The assay was run as described above using 10 µL of the diluted PBT degradation sample.

For the substrate specificity tests, the BHET, MHET and TPA stock solutions were prepared in 100 % DMSO and used to prepare 1:10 dilutions in the activity buffer which then included a final DMSO concentration of 10 % v/v (50 mM NaH_2_PO_4_/Na_2_HPO_4_, 150 mM NaCl, 10 % DMSO, pH 8.0). Serial dilutions were prepared using this buffer condition. The sensor activity assays were performed as described above.

### Circular dichroism (CD) spectroscopy

300 µL of protein samples were prepared with a final concentration of 0.2 mg/mL using activity buffer (50 mM NaH_2_PO_4_/Na_2_HPO_4_, 150 mM NaCl, pH 8.0). CD spectra were obtained on a JASCO J-1500 machine using a glass cuvette with a 1 mm path length. The following settings were used: wavelengths from 240 - 190 nm, a bandwidth of 1 nm, a response time of 2 s, a data pitch of 0.1 nm, a scanning speed 100 nm/min and 5 accumulations. Raw CD data was normalized by subtraction of the respective buffer data and subsequent conversion of the measured signal in millidegree to mean residue molar ellipticity.

### Analytical size exclusion chromatography

500 µL of sensor protein was prepared with a final concentration of 1 mg/mL using activity buffer (50 mM NaH_2_PO_4_/Na_2_HPO_4_, 150 mM NaCl, pH 8.0). The following chromatography was performed on an ÄKTA system. The protein sample was loaded with a 200 µL loop onto a pre-equilibrated Superdex 200 Increase 10/300 GL column followed by 1 CV of activity buffer.

### Isothermal titration calorimetry (ITC)

1 mM TPA stock solution was prepared in activity buffer (50 mM NaH_2_PO_4_/Na_2_HPO_4_, 150 mM NaCl, pH 8.0). All protein and ligand samples were degassed and temperature equilibrated to 25 ℃ for at least 10 min using a degassing station (TA Instruments). 400 µL of protein samples were loaded into the sample cell of a low volume AffinityITC with gold cell (TA Instruments) and ligand solutions were added into the injection syringe (protein and ligand concentrations can be found in Table S2). ITC measurements were performed at 25 ℃ with 20 injections of 2.5 µL (one initial injection of 0.5 µL was excluded from analysis), an adaptive injection interval and stirring rate of 125 rpm. Subtractions of constant heat of dilution values for titration of TPA into activity buffer, peak integration and fitting with a one-site binding model were done with NanoAnalyze (TA Instruments). The reported errors reflect the standard deviation of three technical replicates. Thermograms and binding isotherms are shown in Fig. S3.

### Differential scanning calorimetry (DSC)

Protein concentrations were adjusted to ∼ 3 mg/mL with activity buffer (50 mM NaH_2_PO_4_/Na_2_HPO_4_, 150 mM NaCl, pH 8.0). DSC measurements were performed on a PEAQ DSC (MicroCal) with the following settings: temperature range 20 °C – 110 °C, scanning speed 90 K/h, non-feedback mode. Ligand conditions contained 0.4 mM of TPA dissolved in activity buffer. To establish a thermal history and to allow physical baseline correction, at least two buffer–buffer scans were recorded under identical conditions. The activity buffer served as a reference. Thermograms were normalized to protein concentration. Reversibility of thermal unfolding was assessed by performing a second scan after cooling the samples back to the initial temperature. Data was analyzed and processed using the MicroCal PEAQ DSC software.

### Linker library screening

The library was synthesised and cloned into a pET21b(+)-vector by BioCat GmbH using degenerate codons for the 2 x 2 linker residues. For expression, BL21 (DE3) cells were transformed with the library plasmids and plated on LB agar plates containing 100 µg/mL ampicillin as selection marker. Single colonies were streaked on fresh LB agar plates for colony tracking and subsequently used to inoculate 4 mL of LB media supplemented with 100 µg/mL ampicillin in 24-well blocks. Cells were grown for 6 h at 37 ℃ and 160 rpm. After induction with 1 mM IPTG, protein expression was conducted for 18 h at 20 ℃. Next, the cells were harvested by centrifugation at 4000 rpm for 5 min and the supernatant was removed. Cell pellets were resuspended in 600 µL activity buffer (50 mM NaH_2_PO_4_/Na_2_HPO_4_, 150 mM NaCl, pH 8.0) and transferred to BeadRuptor sample tubes, containing ∼ 125µl of 0.2 mm-sized glass beads. Cell lysis was performed with the BeadRuptor by running 10 cycles of 45 seconds shaking at 5 m/s with 1 minute breaks in between at 4 ℃. Afterwards, cleared cell lysate was obtained by centrifugation for 10 min at 4000 rpm. 40 µL of cleared cell lysate was transferred to a 384-well plate (black, no-bind, Greiner) and either 40 µL of activity buffer or 40 µL of 1 mM TPA (solubilized in activity buffer and pH adjusted) were added. After 5 min of incubation, the fluorescence read-out was obtained using a microplate reader (Spark, Tecan) with the following settings: excitation/emission with monochromator 485/535nm (sf-cpGFP) and 569/592nm (mScarlet3), bandwidth 7.5 nm, mirror: Dichroic 510, gain 60. The dynamic range ΔF/F_0_ was calculated using the TPA-containing sample and the buffer control. To get an estimate of the operational range for variants with ΔF/F_0_ > 1, cleared lysate was titrated with serial dilutions of TPA and using the microplate reader protocol described above. The amino acid sequences of the best variants were obtained by Sanger sequencing.

### Ultra high-performance liquid chromatography (UHPLC)

The procedure was based on a previously reported method ^18^. PBT/PET hydrolysis samples or samples from TPA, MHET or BHET stocks were mixed with acetonitrile consisting of 1 % formic acid in a 1:4 volumetric ratio. Hydrolysis products in the samples were analysed via reversed-phase UHPLC using Thermofisher Ultimate 3000 RS system equipped with a reversed phase C18 column (Kinetex 1.7 μm EVO C18, 100 Å, 50 × 2.1 mm, Phenomenex). A gradient was applied with a flow rate of 1.3 ml/min starting at 100% A (water + 0.1% TFA) and increasing solvent B acetonitrile in the following pattern: 0.04 min 15%, 0.4 min 20%, 0.75 min 50%, 0.95 min 80%, 2.1 min 80%. In total, 1 μl of samples were injected to the column; absorption was measured at 240 nm at a rate of 200 Hz.

### PBT degradation assay

96-well microtiter plates were coated with PBT by adapting the protocol described in Weigert et al. ^18^. The dripping time was reduced to 30 seconds and the drying time was set at exactly 10 minutes.

BL21 (DE3) cells were transformed with a mutant library of LCC-ICCG and plated on LB-Agar plates containing 100 µg/mL ampicillin. Individual clones were transferred to separate wells of a 96-deepwell block containing 1 ml LB-medium supplemented with ampicillin. Precultures were grown overnight at 30 °C. 30 µl were used for inoculating 3 ml Autoinduction medium in 24-deepwell blocks. Cells were grown for 6 - 7 hours at 37 °C and expression occurred overnight at 20 °C. After harvesting, cells were resuspended in 600 µl lysis buffer (50 mM NaH_2_PO_4_/Na_2_HPO_4_, 500 mM NaCl, pH 7.4) and transferred to BeadRuptor sample tubes, containing ∼ 125µl of 0.2 mm-sized glass beads. Cell lysis was performed with the BeadRuptor by running 10 cycles of 45 seconds shaking at 5 m/s with 1 minute breaks in between. Samples were then centrifuged at 4000 x g for 15 minutes and the supernatants were transferred onto 35 µl of Ni^2+^-loaded magnetic beads previously equilibrated in lysis buffer. After incubation at 800 rpm for 1.5 h, the supernatant was discarded and 400 µl wash buffer (50 mM NaH_2_PO_4_/Na_2_HPO_4_, 1 M NaCl, 20 mM imidazole, pH 7.4) was added to the wells and incubated for 15 min. Supernatant was discarded and after an additional wash step with 400 µl borate buffer (50 mM NaCl, 50 mM borate, pH 8.5), 300 µl of borate buffer substituted with TEV-Protease was added and incubated overnight.

Protein concentrations were determined with a Bradford Assay using the Protein Assay Dye from Bio-Rad. 2 parts borate buffer were mixed with 1 part Protein Assay Dye and 60 µl of this mixture were added to individual wells of a black 384-well plate. 40 µl purified protein samples were added per well and mixed by pipetting up and down 5 times. Absorbance at 595 nm was measured after 5 min incubation at room temperature. To calculate individual protein concentrations, a calibration curve was prepared using dilutions of previously purified LCC-ICCG. Protein samples were diluted individually to a concentration of either 200 or 300 nM. 60 µl of these dilutions were then transferred to PBT-coated 96-well microtiter plates and incubated at 80 °C for multiple hours (ranging from 3 - 24 h) and then stored at -20 °C. 10 µl were transferred to a fresh 96-well microtiter plate for dilutions and TPA quantifications with TPAsense. The remaining 50 µl were analysed via UHPLC as described above (Method: Ultra high-performance liquid chromatography (UHPLC).

### PETase and PBTase kinetic measurements

To obtain progress curves of PET and PBT hydrolysis, *Is*PETase or LCC-ICCG was added to PET or PBT-coated 96-well plates with a final concentration of 1 µM dissolved in activity buffer (50 mM NaH_2_PO_4_/Na_2_HPO_4_, 150 mM NaCl, pH 8.0). Each condition was prepared in three technical replicates. The 96-well plates were incubated at 40 ℃ (*Is*PETase) or 80 ℃ (LCC-ICCG) for 2.5 h and samples were taken after 0 h, 0.5 h, 1 h, 1.5 h, 2 h, 2.5 h and stored at 4 ℃ until the TPA quantification.

For the sensor calibration, 100 µL of 400 nM TPAsense3.2 protein (200 nM final) was mixed with 100 µL of either activity buffer or serial dilutions (1:2 ratios) starting from a 200 µM TPA stock in 96-well plates (black, round-bottom, Nunclon, ThermoFisher). After 5 min of incubation, samples were measured in a microplate reader (Spark, Tecan) with the following settings: excitation/emission with monochromator 485/535nm (sf-cpGFP) and 569/592nm (mScarlet3), bandwidth 7.5 nm, mirror: Dichroic 510, gain 80 (sf-cpGFP) and 100 (mScarlet3).

PET and PBT hydrolysis samples were diluted 1:100 in the activity buffer prior to TPA quantification using the protocol as described for the sensor calibration above.

For the ^inv^MM plots, *Is*PETase or LCC-ICCG were added to PET- or PBT-coated 96-well plates in serial dilutions (1:1 ratios) starting with 1 µM (*Is*PETase) and 4 µM (LCC-ICCG) final concentrations dissolved in activity buffer. The 96-well plates were incubated at 20 ℃, 30 ℃, 40 ℃ or 50 ℃ (*Is*PETase) or 80 ℃ (LCC-ICCG) for 2 h (*Is*PETase), 30 min (LCC-ICCG; PBT substrate), and 15 min (LCC-ICCG; PET substrate), respectively. The samples were first transferred to a new 96-well plate and then diluted 1:10, 1:100 or 1:400 (depending on the TPA concentration which was iteratively tested) in the activity buffer prior to TPA quantification using the protocol as described for the sensor calibration above. We estimated a PET and PBT substrate loading of 2.5 g/L per well on the 96-well plates which is subject to a systematic error depending on loss upon dripping and drying while preparing the PET and PBT coatings.

### Preparation of PET particles

PET particles were generated in-house from shredded post-consumer PET bottles (CleanPET WF, Veolia). The material was ground in a Retsch ZM200 centrifugal mill at 18,000 rpm using a 4,000 µm spacer sieve (perforated sieve) with a 12-tooth rotor. The fractionation related to particle size was performed in an air jet sieve (e200 LS, Hosokawa Alpine) using pore sizes of 4000, 3150, 2000, 600, and 250 µm.

### PET microplastic detection in wastewater

Wastewater samples were taken from the wastewater treatment plant in Bayreuth on 19.11.2025 at around 2 pm. Sample locations are indicated in Fig. 4a.

To test the activity of TPAsense3.2 in wastewater samples, small particles were removed by filtering 1 mL of each sample with a 0.22 µm cut-off filter. Then, serial dilutions of TPA were prepared in each wastewater sample using a 1 mM TPA stock solution solubilized in the activity buffer (50 mM NaH_2_PO_4_/Na_2_HPO_4_, 150 mM NaCl, pH 8.0). Serial dilutions of TPA in the activity buffer were used as a control. The sensor activity assay was performed as described above (see Method: Sensor activity assay in vitro). During data analysis, we observed that the highest TPA concentration showed a strongly reduced signal ratio to the next following dilutions which might be due to the higher content of activity buffer in these samples. That is why the highest concentration was removed from the analysis. One value of the influent calibration showed an unusually low mScarlet3 fluorescence intensity value indicating a pipetting error and was thus removed from further analysis.

For the PET microplastic detection experiments, a final concentration of 500 nM of the LCC-ICCG was added to 1 mL of unfiltered wastewater samples. The positive control was spiked with one small spatula of < 250 µm PET particles (see Method: Preparation of PET particles). The negative control received neither enzyme nor PET particles. All samples were prepared in triplicates. They were incubated at 70 ℃ for 24 h with 200 rpm shaking. Afterwards, remaining particles were removed by centrifugation 15,000 x g for 10 min (Eppendorf 5427R). The sensor activity assay was performed as described above (see Method: Sensor activity assay in vitro).

In general, the amount of TPA released from PET microplastic in wastewater samples could be quantified but it is not clear if PET was degraded to completion after 24 h. For this, control measurements of samples spiked with defined amounts of PET would be required ^41^.

### Data analysis

The sensor activity data obtained by in vitro titration experiments was fit to a scaled version of the Hill-Langmuir equation:

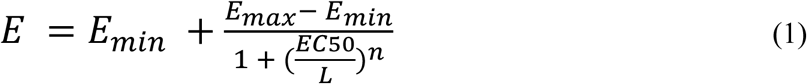

with *E* as the biosensor output signal, *E_min_* and *E_max_* as the baseline and maximal signal, respectively, *L* is the ligand concentration in molar units, *EC50* is the half-maximal effective concentration, and *n* is the Hill coefficient. To calculate the ligand concentration from the biosensor output signal, the Hill-equation was solved for *L*:

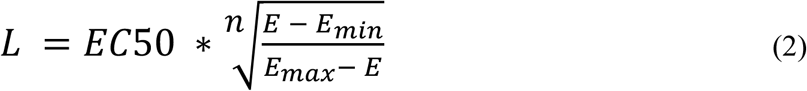

and values for the parameters were taken from the calibration (fitting data to equation 1). For the analysis of PETase and PBTase kinetics, the inverse Michaelis-Menten equation was used to fit the experimental data as discussed in Kari et.al. ^35^:

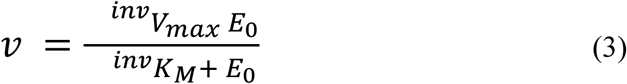

with *v* as the reaction velocity in units of nmol s^-1^ g^-1^, *E_0_* as the initial enzyme concentration, *^inv^V_max_* as the maximal velocity when the substrate is fully saturated with enzyme and ^inv^*K_M_* (inverse Michaelis-Menten constant) as the enzyme concentration when the half-maximal velocity is reached. If not stated otherwise, all data analysis was done with python.

## Acknowledgements

We thank Sabrina Wischt and Tristan Mühlbauer for experimental support and SFB 1357 Microplastics subproject Z01 for production of PET microplastic particles. This study was supported by the Deutsche Forschungsgemeinschaft (DFG, German Research Foundation—Project Number 391977956—SFB 1357 Microplastics, subproject C03) and the European Union’s Horizon 2020 research and innovation program under Marie Skłodowska-Curie Grant Agreement 955623. AG and BH acknowledge BASF SE (Ludwigshafen am Rhein, Germany) for generous funding and providing PBT.

## Supporting information

**Supporting Table 1.**
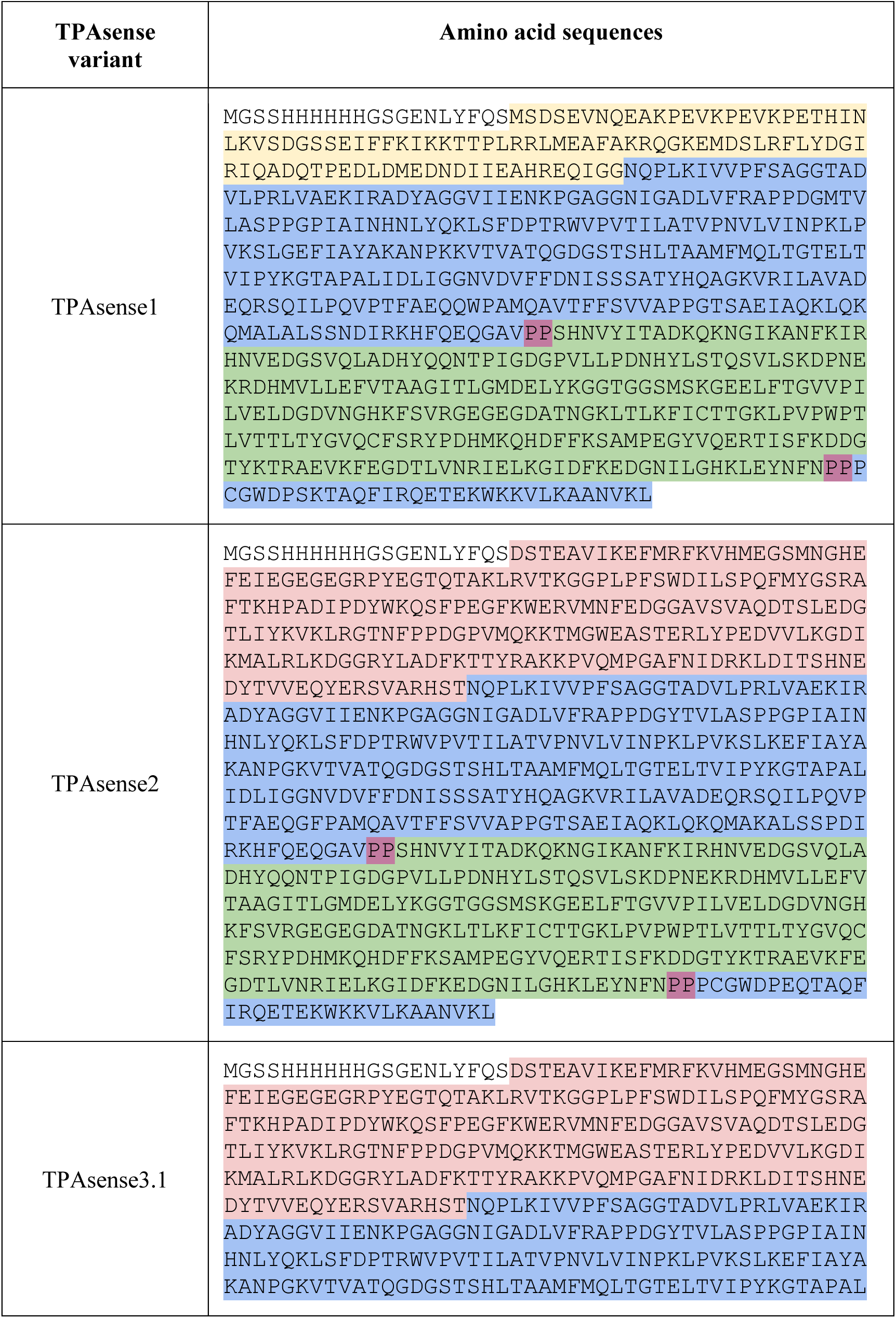
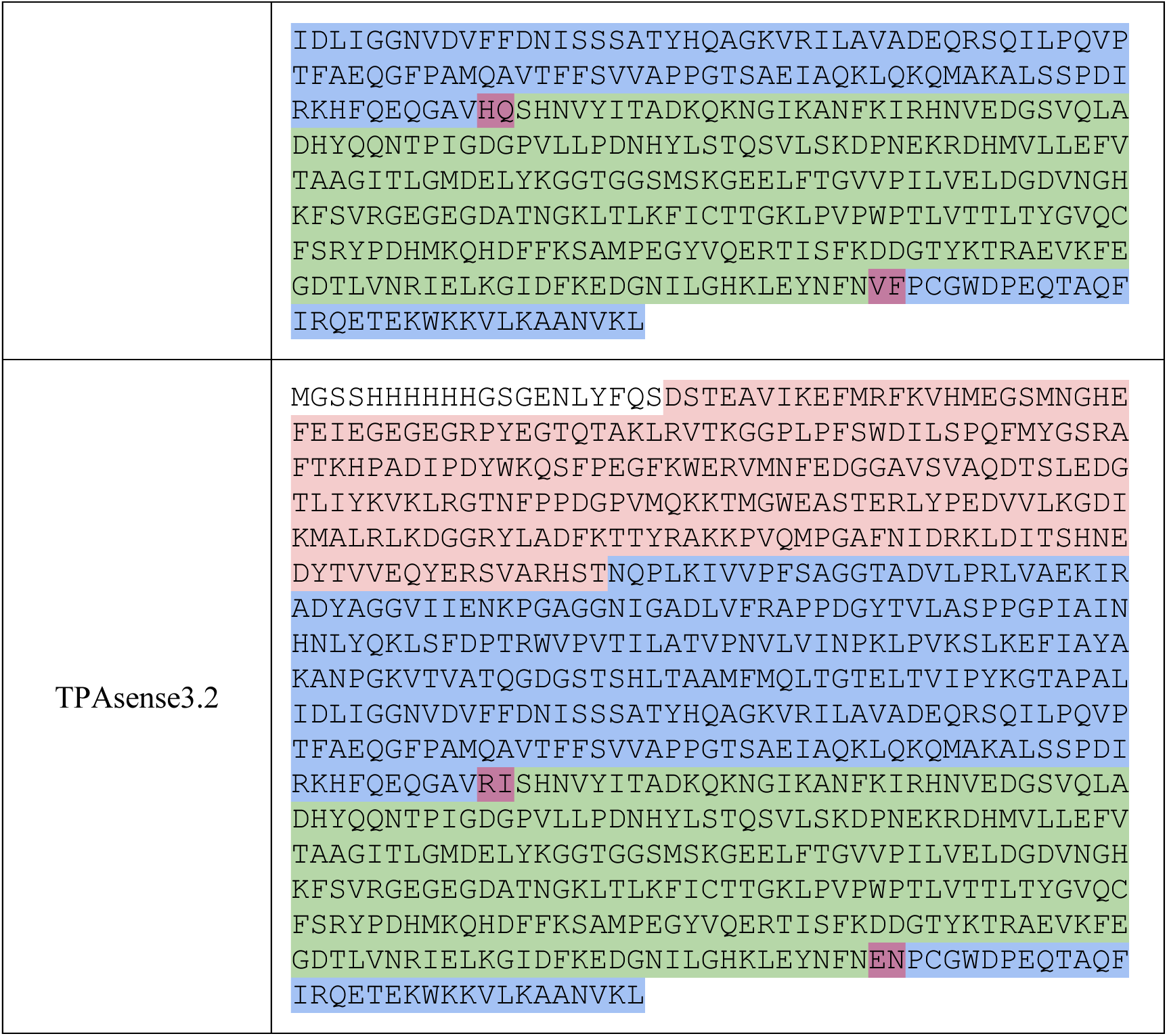
Amino acid sequence of TPAsense variants. SUMO tag (yellow), TphC (blue), linker residues (purple), sf-cpGFP (green) and mScarlet3 (red) are marked in color.

**Supporting Table 2.**
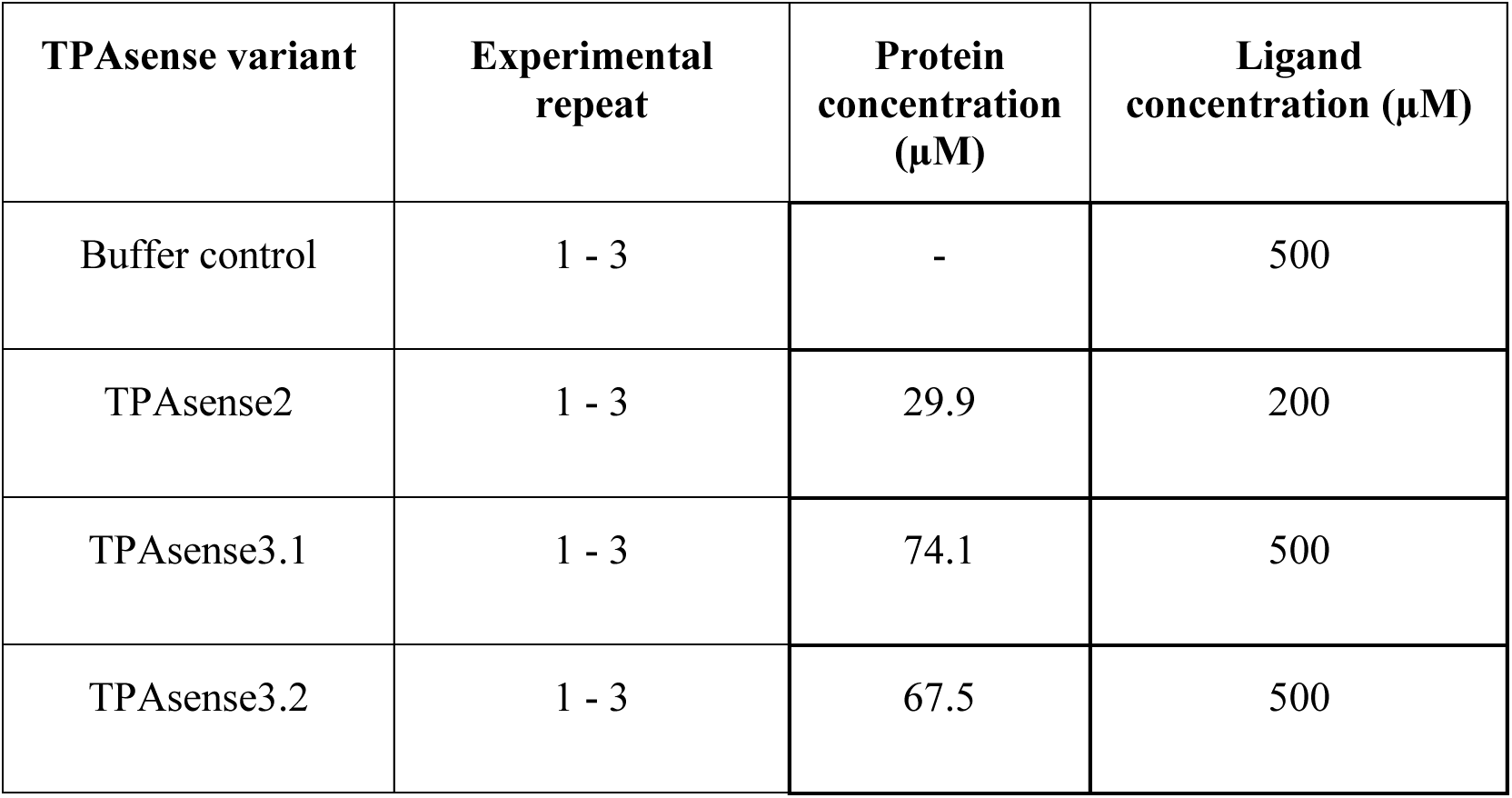
Protein and ligand concentrations used in ITC experiments.

**Supporting Table 3.**
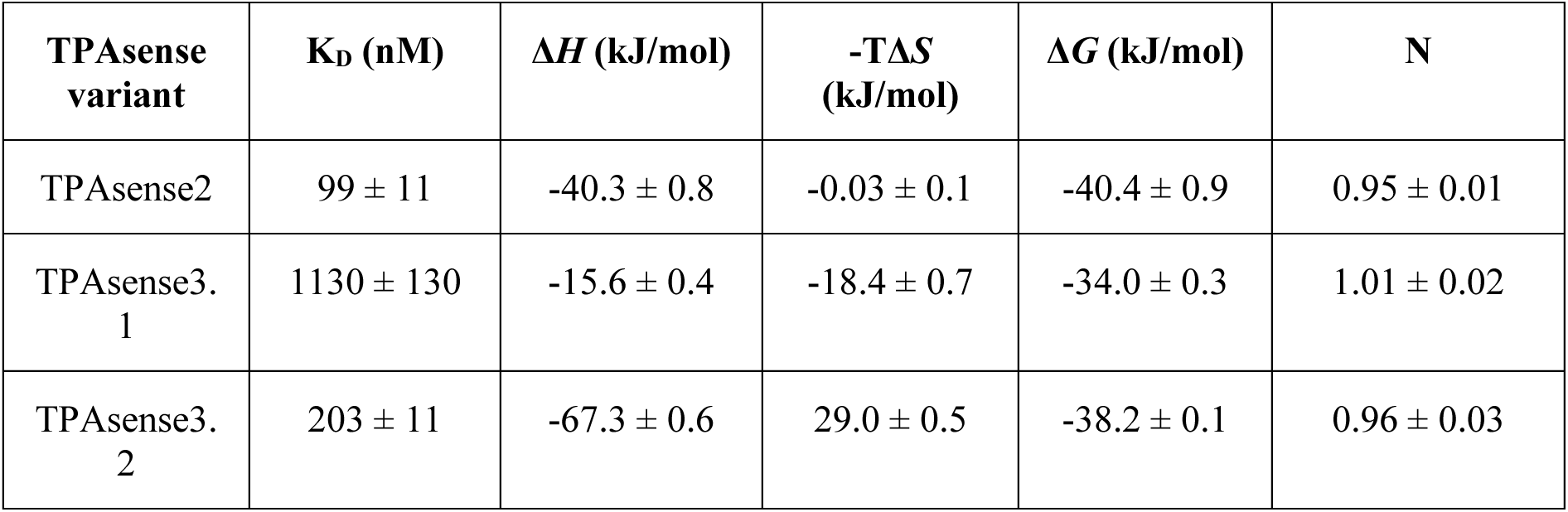
Thermodynamic parameters of TPAsense2 and TPAsense3 variants binding to TPA measured by ITC. Averages and standard deviations were calculated based on three technical repeats.

**Supporting Figure 1.**
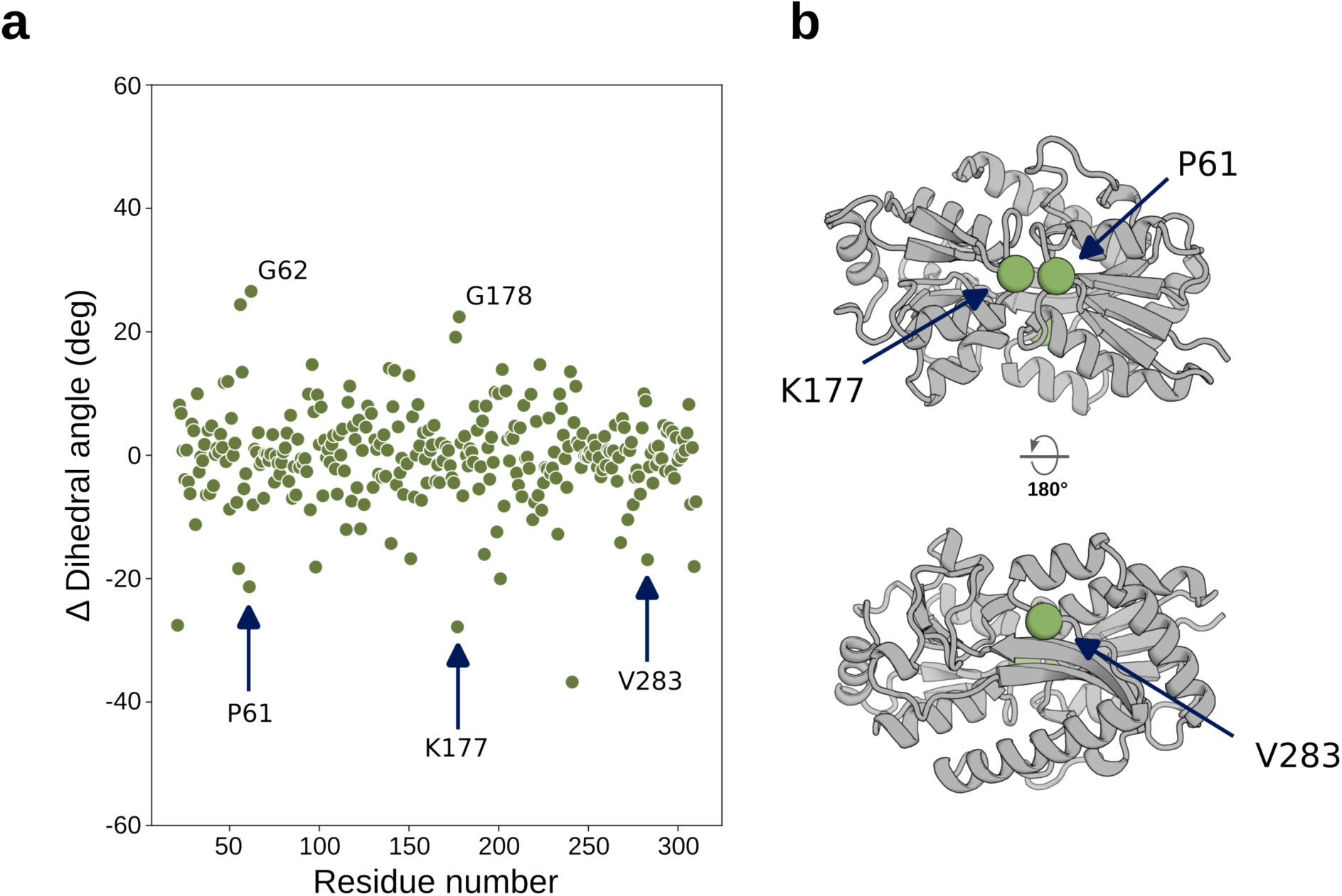
Changes in backbone dihedrals revealed potential insertion sites. **a**, Dihedral angle changes of the backbone of TphC open and closed states. Insertion sites tested in this study are marked with arrows. **b**, Closed state crystal structure of TphC (PDB ID: 7NDS). Residues after which sf-cpGFP was inserted are marked as green spheres.

**Supporting Figure 2.**
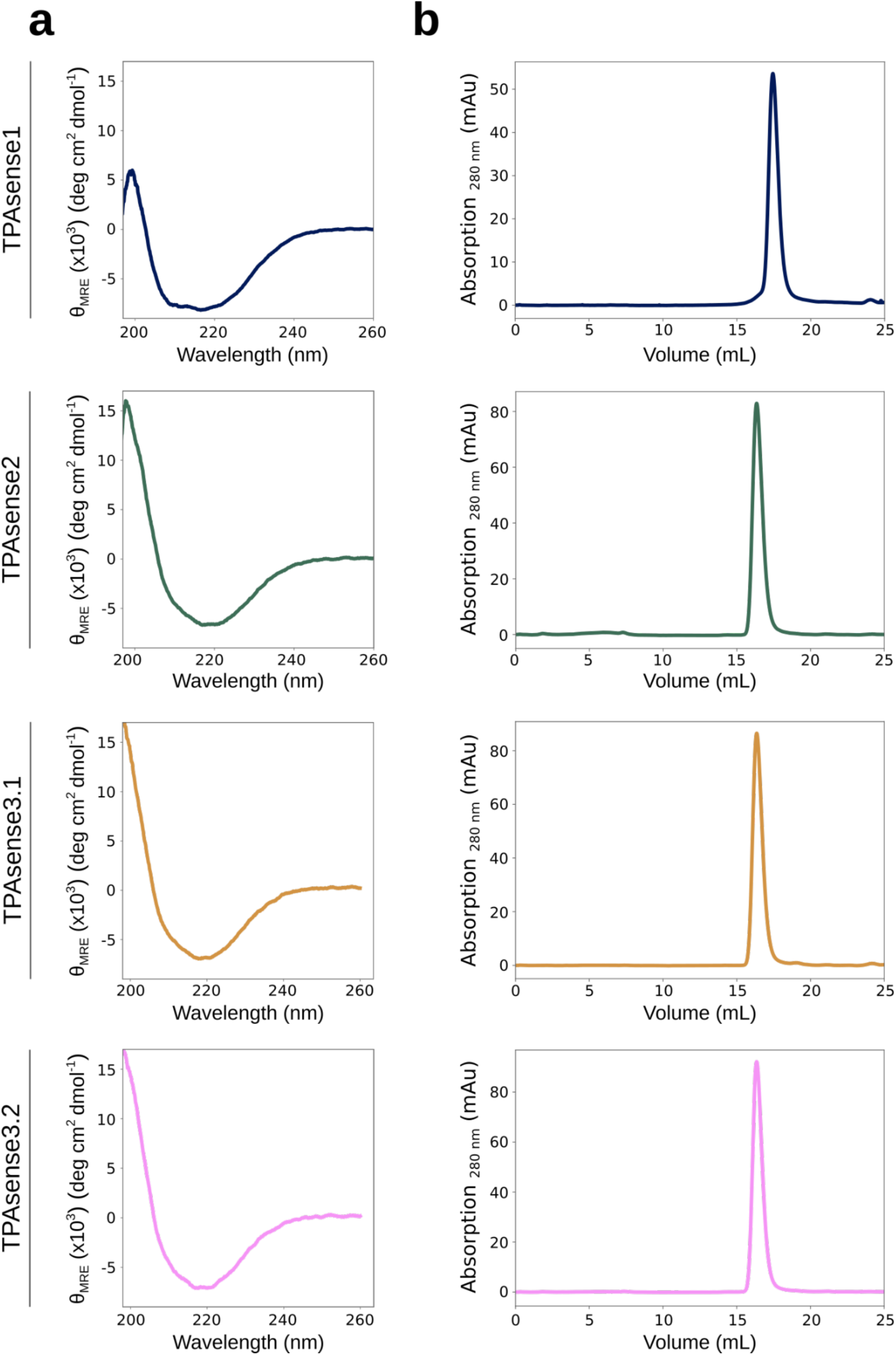
All TPAsense variants were properly structured and free of aggregation or formation of multimeric species. **a**, Circular dichroism (CD) spectra plotted as mean-residual ellipticity from 198-260 nm. **b**, Analytical SEC chromatograms of TPAsense variants followed by absorption at 280 nm.

**Supporting Figure 3.**
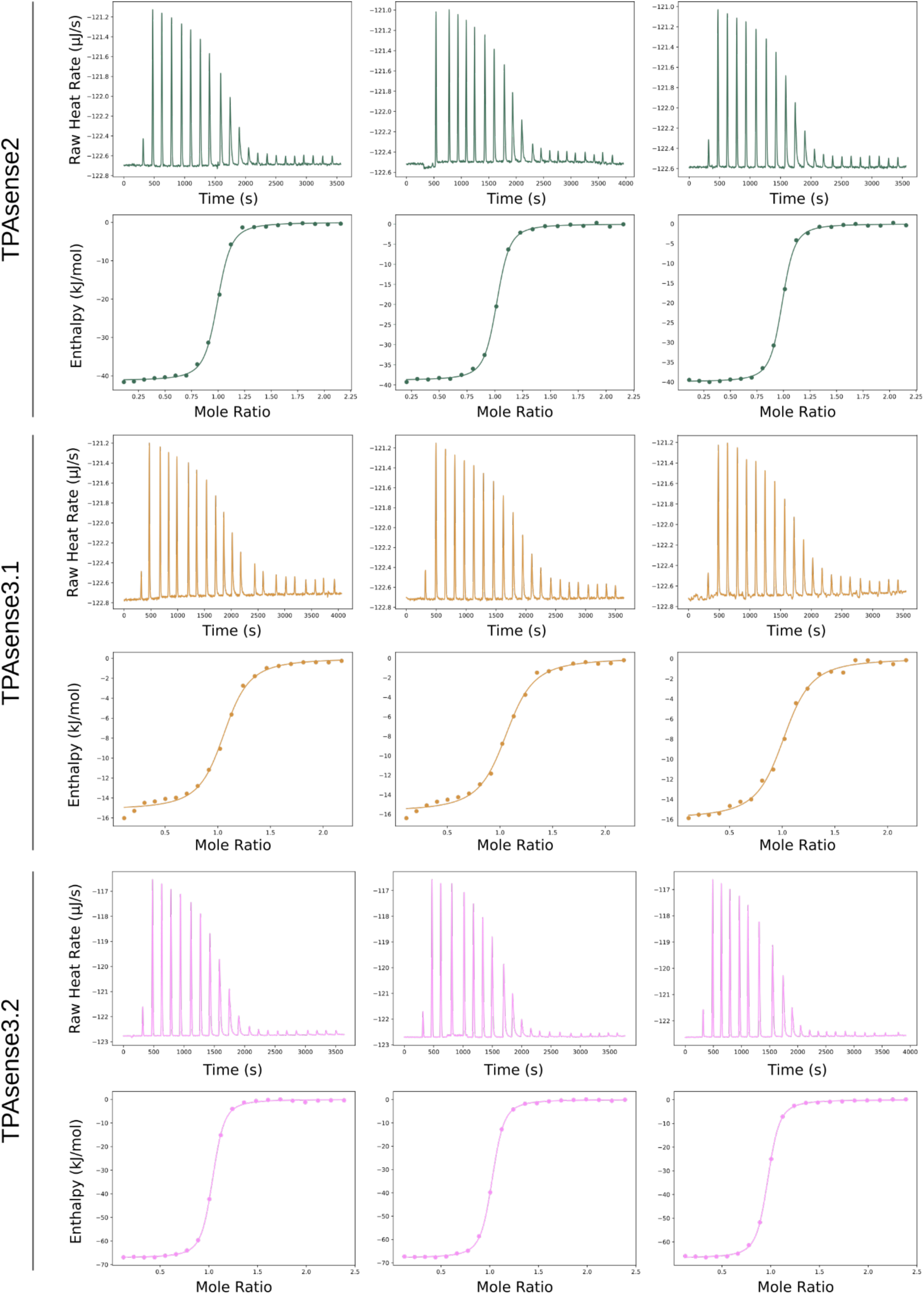
Thermograms and binding isotherms of TPAsense2 and TPAsense3 variants with terephthalate in three technical repeats. Buffer-terephthalate control titrations were performed to obtain baseline values that were subtracted from the integrated heats of TPAsense2 and TPAsense3 variants.

**Supporting Figure 4.**
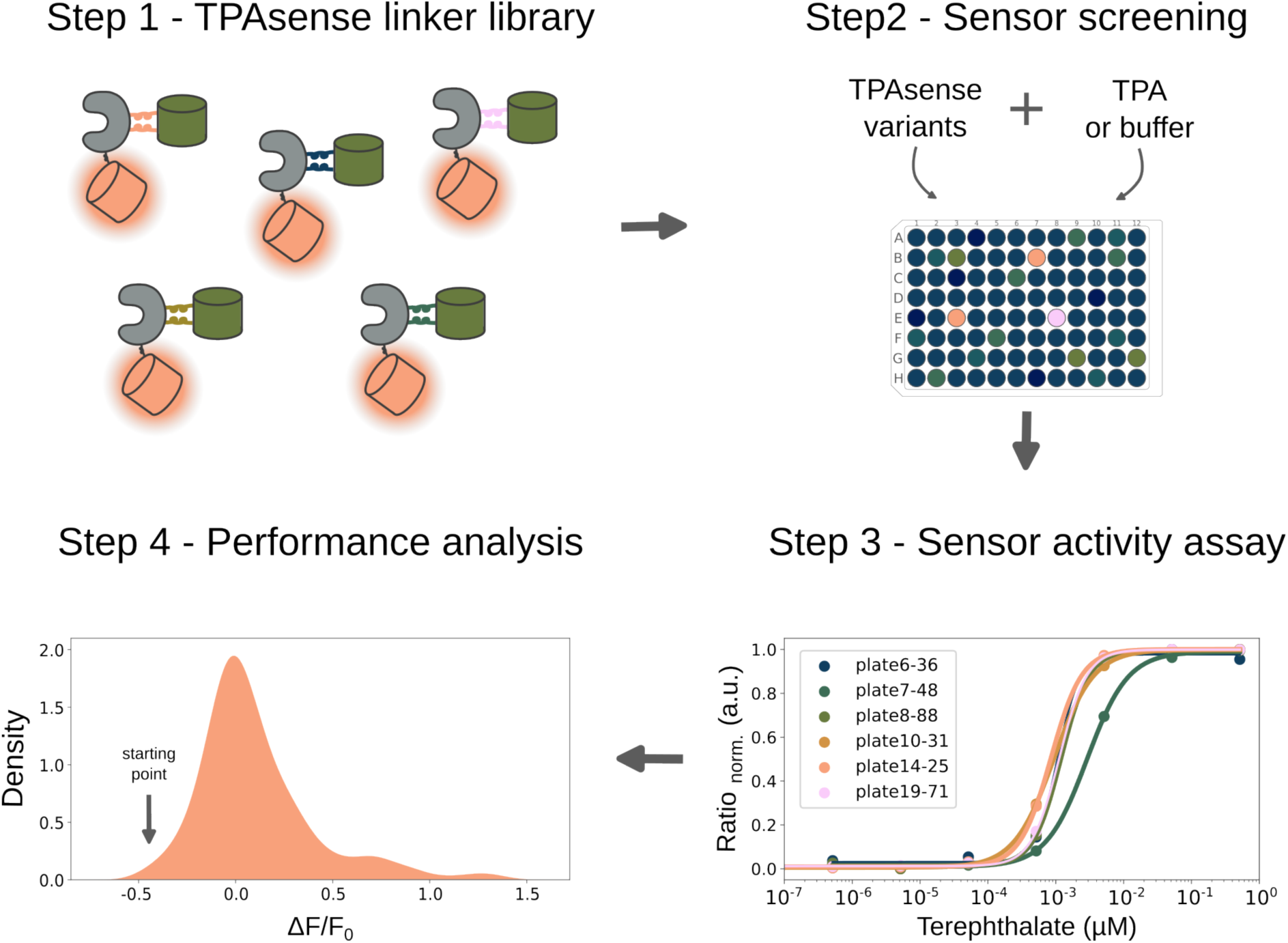
TPAsense linker library screening workflow in lysate. First, *E.coli* were transformed with a TPAsense library, clones were picked and cultivated for expression in 24 deep well plates (step 1). After cell lysis, TPAsense variants were screened by addition of TPA compared to buffer controls in 384 well format (step 2). The best variants were titrated with a serial dilution of TPA in lysate to obtain estimates for the dynamic and operational range (step 3). Finally, the screening results of 570 variants were analyzed (step 4). It turned out that the initial variant, TPAsense2, exhibited the most negative ΔF/F_0_ (arrow in step 4). Most linker variants did not show a TPA-dependent signal change and only a very small portion showed ΔF/F_0_ values > 1.

**Supporting Figure 5.**
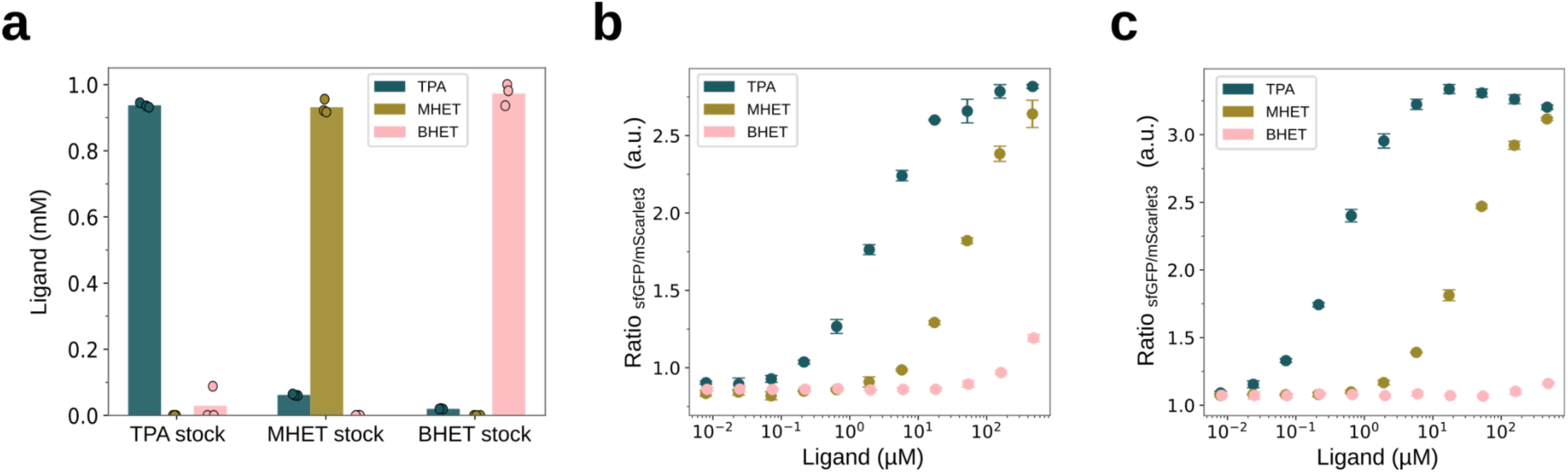
Substrate specificity of TPAsense3 variants. **a**, UHPLC analysis of 1 mM TPA, MHET and BHET stock solutions. Presence of BHET in the TPA stock was only present in one out of three technical repeats and can thus be considered an outlier. **b**, In vitro titrations of TPAsense3.1 with serial dilutions of TPA, MHET and BHET. **c**, In vitro titrations of TPAsense3.2 with serial dilutions of TPA, MHET and BHET.

**Supporting Figure 6.**
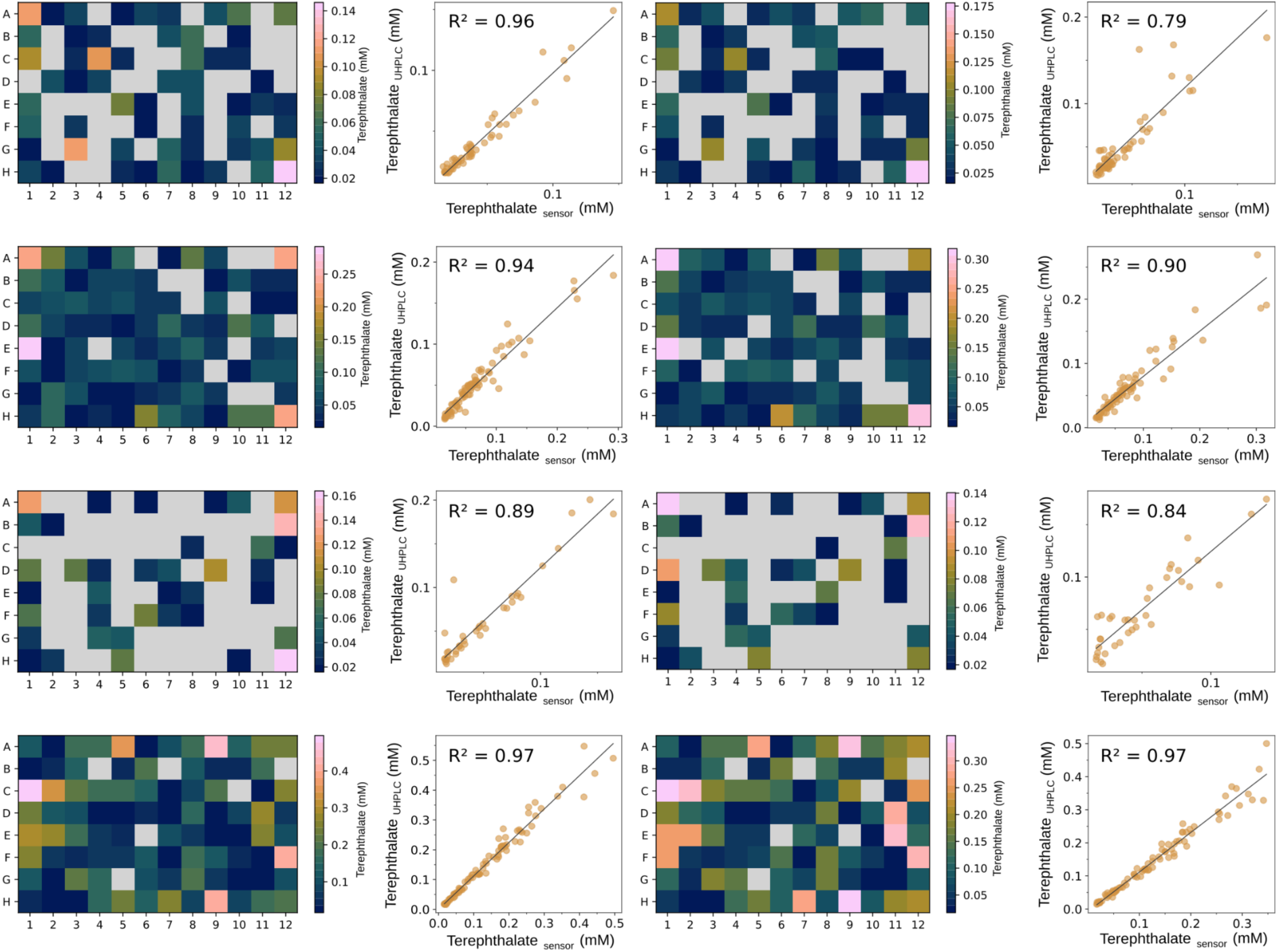
Screening PBT degradation of a LCC mutant library with TPAsense3.1 for all 96 well plates and comparison to UHPLC data. Values outside the boundaries of the operational range of TPAsense3.1 were filtered out (gray boxes). The TPA titres determined with TPAsense3.1 were compared to those measured with UHPLC and the coefficient of determination R^2^ was calculated using a linear regression model.

**Supporting Figure 7.**
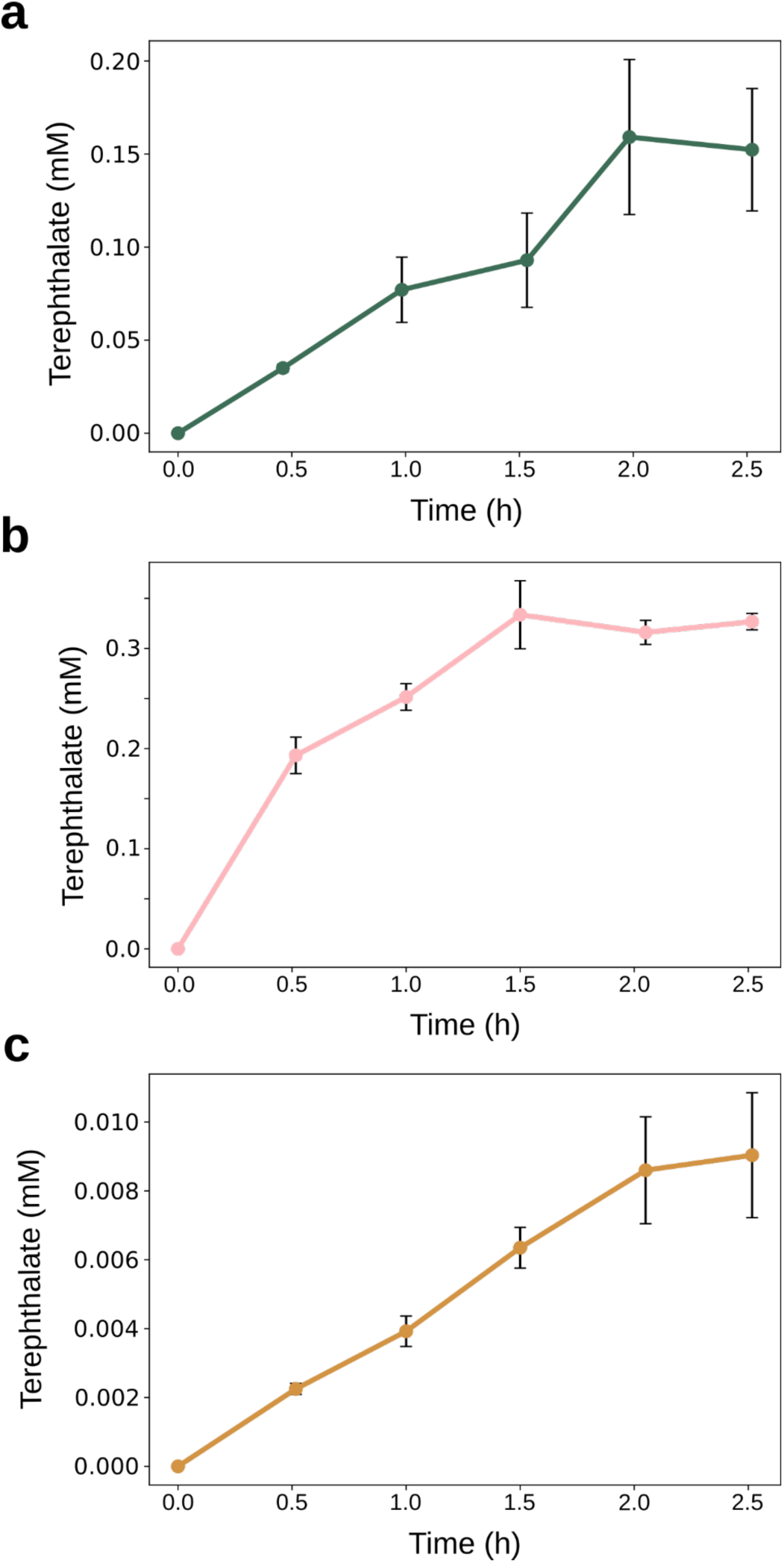
Progress curves of enzymatic PET and PBT hydrolysis. Reactions were followed over 2.5 h with samples taken every 30 min. Data points and error bars reflect averages and standard deviations of three technical repeats, respectively. **a**, PET hydrolysis with IsPETase at 40 ℃. Based on this curve, the reactions for the ^inv^MM approach were stopped after 2 h. **b**, PET hydrolysis with LCC-ICCG at 80 ℃. Based on this curve, the reactions for the ^inv^MM approach were stopped after 15 min. **c**, PBT hydrolysis with LCC-ICCG at 80 ℃. Based on this curve, the reactions for the ^inv^MM approach were stopped after 30 min.

